# Inertial Focusing of Particles and Cells in the Microfluidic Labyrinth Device: Role of Sharp Turns

**DOI:** 10.1101/2022.06.02.494575

**Authors:** Anirudh Gangadhar, Siva A. Vanapalli

## Abstract

Inertial, size-based focusing was investigated in the microfluidic labyrinth device consisting of several U-shaped turns along with circular loops. Turns are associated with tight curvature, and therefore induce strong Dean forces for separating particles, however, systematic studies exploring this possibility do not exist. We characterized the focusing dynamics of different-sized rigid particles, cancer cells and white blood cells over a range of fluid Reynolds numbers *Re*_*f*_. Streak widths of the focused particle streams at all the turns showed intermittent fluctuations which were substantial for smaller particles and at higher *Re*_*f*_. In contrast, cell streaks were less prone to fluctuations. Computational fluid dynamics simulations revealed the existence of strong turn-induced Dean vortices which help explain the intermittent fluctuations seen in particle focusing. Next, we developed a measure of pairwise separability to evaluate the quality of separation between focused streams of two different particle sizes. Using this, we assessed the impact of a single sharp turn on separation. In general, the separability was found to vary significantly as particles traversed the tight-curvature U-turn. Comparing the separability at the entry and exit sections, we found that turns either improved or reduced separation between different-sized particles depending on *Re*_*f*_. Finally, we evaluated the separability at the downstream expansion section to quantify the performance of the labyrinth device in terms of achieving size-based enrichment of particles and cells. Overall, our results show that turns are better for cell focusing and separation given that they are more immune to curvature-driven fluctuations in comparison to rigid particles.

## 1. Introduction

Inertial microfluidics has gained prominence in a variety of applications such as sheath-less flow cytometry ^1-7^, fast particle transfer across fluid streams ^3^, filtration of biological matter ^6,8,9^ and separating cancer cells from blood cells ^10,11^. In one manifestation of inertial microfluidics, curved geometries are used, where, in addition to inertial forces, strong curvature forces driven by centrifugal effects are introduced ^12^. These so-called Dean flows can manipulate differential migration of particles or cells, focusing them at different lateral positions depending on their sizes ^13^. Such inertial microfluidic systems incorporating channel curvature have been shown to be efficient at processing large volumes of suspensions in a relatively short time with good rates of particle separation efficacy and purity ^10,11^.

In straight channels, particles with finite inertia get focused due to a balance of shear gradient lift force and the viscous Stokes drag ^14^. However, in curved channels, the fluid is pushed radially outwards due to centrifugal forces which sets up a transverse pressure gradient. Thus, fluid near the top and bottom walls moves inward resulting in two symmetric, counter-rotating cross-sectional vortices or Dean vortices ^15,16^. The onset of these vortices is determined by the Dean number^12^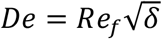, where *Re*_*f*_ is the fluid Reynolds number and *δ* is the curvature ratio. In addition to the aforementioned forces in straight channels, this vortex gives rise to an additional force called the Dean drag force acting on particles flowing in curved geometries ^12^. Here, *Re*_*f*_ = *ρ*_*f*_*U*_*f*_*D*_*h*_/*μ*_*f*_ is the fluid Reynolds number (*ρ*_*f*_ and *μ*_*f*_ denote fluid density and dynamic viscosity respectively, *U*_*f*_ is the mean fluid velocity in the channel and *D*_*h*_ is the hydraulic diameter) and the curvature ratio *δ* = *D*_*h*_/2*R*, where *R* is the mean channel radius of curvature.

The basic mechanism for particle separation in curved channels appears to be that the inertial lift force *F*_*L*_ stabilizes particle position (i.e., particle focusing), while the Dean drag force *F*_*D*_ aids in lateral migration due to the cross-sectional circulation (i.e., particle separation) ^12,17^. Both these forces are dependent on particle diameter *d*_*p*_, the fluid Reynolds number *Re*_*f*_ and channel curvature ratio *δ* requiring optimization of channel geometries and flow conditions to achieve high fidelity in focusing and particle separation ^12^. Experimentally, two geometrical parameters are available to optimize particle focusing and separation in curved channels. The first is the focusing length *L*_*f*_, which is defined as the channel length required to focus particles. Since *L*_*f*_ depends strongly on particle diameter *d*_*p*_, i.e. 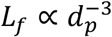, larger particles can be focused at short channel lengths^12^. Smaller particles in contrast require much longer channel lengths to focus due to the reduced inertial lift forces *F*_*L*_ acting on them since 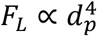. The second is the curvature ratio, where tighter curvatures lead to higher Dean drag forces. Since 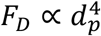, Dean drag forces are weaker for smaller particles, necessitating tighter curvatures to focus them.

An important application where size-based separation in curved geometries has been effective is the separation of cancer cells from blood cells in the context of isolating circulating tumor cells from patient blood ^18-21^. The diameter of tumor cells, white blood cells (WBCs) and red blood cells (RBCs) lie in the range of ∼10-20 *μm*, ∼7-12 *μm* and ∼8 *μm* respectively ^18^. Given these differences in cell size, spiral geometries where Dean flow can be invoked have been developed for separating cancer cells ^18-21^. However, a potential drawback of the spiral geometries is that smaller cells or particles might be difficult to focus due to lack of tight curvatures that can induce strong mixing forces.

Recently, a microfluidic labyrinth device was proposed that isolates cancer cells from blood cells ^22^, with the aim of improving the focusing of smaller-sized blood cells. In addition to spiral sections, the labyrinth device contains multiple sharp turns. The turns were proposed to help in two ways: (i) allow longer channel lengths to be packaged into a small device footprint, thereby increasing the opportunity to focus smaller particles and (ii) augment the local Dean forces due to their tight curvatures which helps to push smaller particles towards their equilibrium positions, focusing them. Another study by ^23^ also suggested that turns can enhance the focusing of smaller particles. Despite the proposed hypothesis that turns can aid in focusing of smaller particles and therefore provide efficient separation, currently quantitative data characterizing how particle size and fluid Reynolds number influence the focusing dynamics is lacking. Moreover, the turns present are U-shaped which leads to quick and successive changes in flow direction that can complicate focusing dynamics, which is yet to be quantified.

In this study, we use the labyrinth device as a prototypical system that contains both spiral arcs and turns to investigate how particle size and fluid Reynolds number influence the focusing dynamics at all the corners present in the device. Additionally, we simulate the fluid flow in the U-turn region to understand how variations in focusing dynamics can be explained by the cross-sectional flow fields. Next, we develop a quantitative measure of separability S to track the separation between focused streams of size-differing particles as they approach and exit the sharp turn. A comparison of S before and after the turn allows us to determine whether the tight-curvature turns helps to improve particle separation or not and if so, at what flow conditions. Finally, we characterize the separability in the downstream expansion section where particle streams are collected, to report on the overall performance of the labyrinth device for size-based separation of particles and cells.

## 2. Experimental methods

### 2.1 Labyrinth device design

The microfluidic labyrinth device has a width of 500 *μm* and a total length of 637 mm. It consists of 11 spiral loops and 56 corners, with the curvature ratio in the loops varying from 5.29 × 10^−3^ to 3.7 × 10^−2^. The channel height *H* is uniformly 97 um. Multiple sharp turns throughout the device induce an abrupt increase in geometrical curvature. The fabricated devices were received from the Sunitha Nagrath laboratory at the University of Michigan.

### 2.2 Sample preparation

In this study, three types of samples were used: MCF-7 breast cancer cells, WBCs and fluorescent polystyrene microspheres. MCF-7 breast cancer cells were cultured and stained with a live fluorescent dye (CellTracker™ Red CMTPX dye, Invitrogen™, Carlsbad, USA). WBCs were isolated from human whole blood and also stained with a live cell marker (10 *μM* CMTPX, Invitrogen™, Carlsbad, USA). Detailed methods for MCF-7 cell culture and staining of cancer cells and WBCs are provided in our previous works ^24-26^. Tagged WBCs and MCF-7 cells with mean size 10, 21 *μm* respectively were diluted with 1X phosphate buffer saline to make up working concentrations of 100,000 and 500,000 cells/ml. Gentle pipetting prior to the experiment ensured that the cells did not aggregate or settle.

Fluorescent polystyrene beads of four different sizes were selected with sizes comparable to the cell types used in the study. 7.32 *μm* TRITC (Bangs Laboratories Inc., Fishers, USA), 12 *μm* FITC, 15 *μm* TRITC (Thermo Fisher, Carlsbad, USA) and 20 *μm* FITC (Polysciences Inc., Warrington, USA) particles were suspended in deionized water to make up a final concentration of 1 × 10^6^ beads/ml. 0.1% (V/V) Tween 20 (Sigma-Aldrich, St. Louis, USA) was added to minimize aggregation of microspheres. Particle suspensions were vortexed prior to the experiment to ensure adequate dispersion of the microspheres in suspension.

### 2.3 Flow conditions

Samples were processed through the devices using a syringe pump (PHD 2000, Harvard Apparatus, Holliston, USA). Prior to the experiment, devices were flowed with 1% Pluronic^®^ F-127 (Sigma-Aldrich, St. Louis, USA) solution, diluted in 1X PBS at 100 *μL*/*min* for 10 minutes followed by incubation at room temperature for 30 minutes to minimize unspecific cell adhesion to microchannel walls ^22^. In order to experimentally vary the fluid Reynolds number *Re*_*f*_, four different flow rates were tested: 1.5, 2.5, 3.5 and 5 ml/min. Table 1 shows the experimental parameter space explored in this study.

**Table 1:** Range of dimensionless parameters used in the study.

|  |  |
| --- | --- |
| Confinement ratio, $\lambda$ | 0.04 - 0.13 |
| Fluid Reynolds number, $Re_f$ | 83, 139, 194, 278 |
| Particle Reynolds number, $Re_p$ | 0.16 - 4.41 |
| Dean number, $De$ | 6 - 53 |

### 2.4 Fluorescent streakline imaging and streak width measurement

Imaging was performed using an Olympus IX81 microscope (Massachusetts, USA). After 1 minute of flow stabilization, fluorescence streak images were recorded using a Hamamatsu digital camera (ImagEM X2 EM-CCD, New Jersey, USA). At each flow rate tested, an automated stage (Thorlabs, New Jersey, USA) was used to capture images at all 56 corners and expansion region of the device at multiple time points using the Slidebook 6.1 software (3i Intelligent Imaging Innovations Inc., Denver, USA). Images were taken at x4 objective magnification with an exposure time of 100 ms and frame rate of 10 fps. ImageJ software (NIH, Bethesda, USA) was used to obtain a single, mean intensity image from a stack of images at every corner.

Streak width was obtained using ImageJ software (NIH, Bethesda, USA). At the corners, this was measured as the width of the fluorescence streak (in micron) along the line connecting the inner and outer corner vertices. Previous attempts to standardize our measurements involved taking a line scan connecting the two corner vertices and calculating the full width at half-maximum (FWHM) of the Gaussian-fit intensity profile ^27^. However, it was discovered that the fit was below par in many cases. We attribute this to the strongly asymmetric nature of the corner geometry. Moreover, tight curvatures at the turns strongly impact the particle streaks and, in many cases, they were found to deviate significantly from the classical Gaussian intensity profile. Subsequently, we decided to measure the streak width using the above-mentioned method.

### 2.5 Computational fluid dynamics simulations

To visualize flow near the corner, 3D CFD fluid simulations were performed using ANSYS Fluent (v. 19.3). The geometry was imported from AutoCAD (v. 2019, Autodesk). We selected a region containing corners 55 and 56 of the Labyrinth device. Steady-state viscous laminar model was chosen with water as the working fluid. No slip condition was applied at the microchannel walls. At the inlet, a velocity boundary condition was imposed to match the experimental flow rates while an outflow condition was imposed at the outlet. For discretization, we chose a mesh resolution of 20 *μm* in the streamwise plane and 10 *μm* for the cross-section. To solve for the flow field, SIMPLEC algorithm was selected for pressure-velocity coupling along with least squares-based gradient and second order scheme for pressure and momentum. The maximum number of iterations was set to 1000 and convergence tolerance was 10^−6^. For computing the 2D cross-sectional flow streamlines, the number of spatial points was set to 300.

## 3. Results and discussion

### 3.1 Dynamics of particle focusing at the turns in the labyrinth device

The geometry of the microfluidic labyrinth device is complex compared to standard curved or spiral geometries. Two important differences are (i) the existence of multiple U-shaped turns in the labyrinth compared to curved geometries (Fig. 1b), where the turns might induce strong circulatory forces affecting particle focusing (ii) unlike spiral geometries, where, typically, the curvature ratio *δ* monotonically decreases from inlet to outlet due to increasing channel radius of curvature, the curvature ratio is non-monotonic in the labyrinth (Fig. 1c). Both the U-shaped turns and the non-monotonic curvature ratio may lead to variation in focusing as particles traverse through the labyrinth. Thus, we programmed the microscope stage to image particle focusing in the corner locations (Fig. 1d), enabling us to understand the role that sharp turns play on determining the particle focusing behavior.

**Figure 1.**
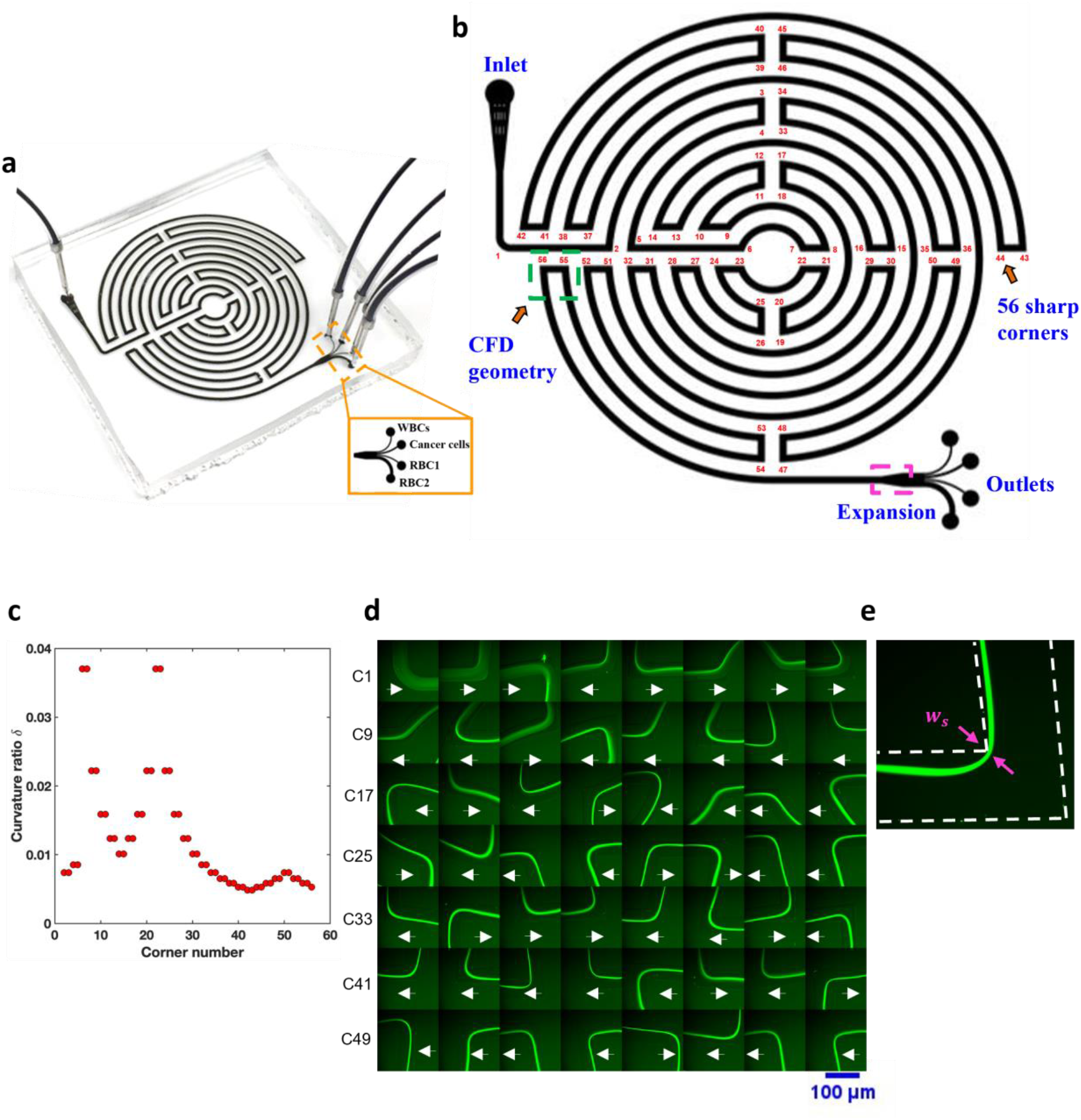
Labyrinth device geometry and characterization of particle focusing at the turns. (a) Microfluidic labyrinth device containing 4 separation outlets to collect WBCs, cancer cells and RBCs; (b) 2D geometry of labyrinth highlighting the location of corners, numbered 1-56 as well as the expansion region and separation outlets. Geometry used for CFD simulations is indicated by the green rectangle; (c) Variation of curvature ratio *δ* plotted against the corner number preceding the curved channel; (d) Fluorescent streak images of 12 *μm* polystyrene microspheres at all 56 corners of the labyrinth device, displayed as a montage, *Re*_*f*_ = 83. Arrows indicate direction of the flow; (e) Width of the fluorescent particle streak *w*_*s*_ is measured along the line joining the two vertices associated with the corner.

To characterize particle focusing at the turns of the labyrinth device, we varied particle diameter *d*_*p*_ and the fluid Reynolds number *Re*_*f*_ and quantified the dispersity in particle focusing. Polystyrene microspheres of four different sizes: 7, 12, 15 and 20 *μm* as well as MCF-7 breast cancer cells, WBCs and RBCs were tested at four different flow rates: 1.5, 2.5, 3.5 and 5 *ml*/*min* corresponding to *Re*_*f*_ = 83, 139, 194, 278 respectively. From the acquired fluorescence streak images (Fig. 1d), the streak width *w*_*s*_ was measured at every corner (Fig. 1e). To assess the experimental variability in the streak width measurement, we performed three independent trials with the 7 and 12 *μm* particles at *Re*_*f*_ = 139, and evaluated the coefficient of variation (CoV) at each of the 56 corners. We found the mean CoV from the 56 corners to be within 9 and 12 % respectively for 7 and 12 *μm* particles at *Re*_*f*_ = 139 (Fig. S1), suggesting good reproducibility in streak width measurements.

Fig. 2 shows the streak widths at all the labyrinth corners for all the conditions investigated. In all cases, even though the general trend is that the streak width declines with corner number, we observe that the streak width data shows intermittent fluctuations. Stated differently, even though particles appear to focus at specific locations evident by the lower streak width values, they tend to de-focus at other locations as they traverse through the multiple sharp turns in the labyrinth device. Qualitatively, the degree of these intermittent fluctuations is high for smaller particles and at higher Reynolds numbers. Interestingly, the MCF-7 and WBC cells show the least fluctuations at the lower Reynolds numbers. We note that WBCs at *Re*_*f*_ = 278 and the RBCs could not focus, so therefore streak width data is not shown.

**Figure 2.**
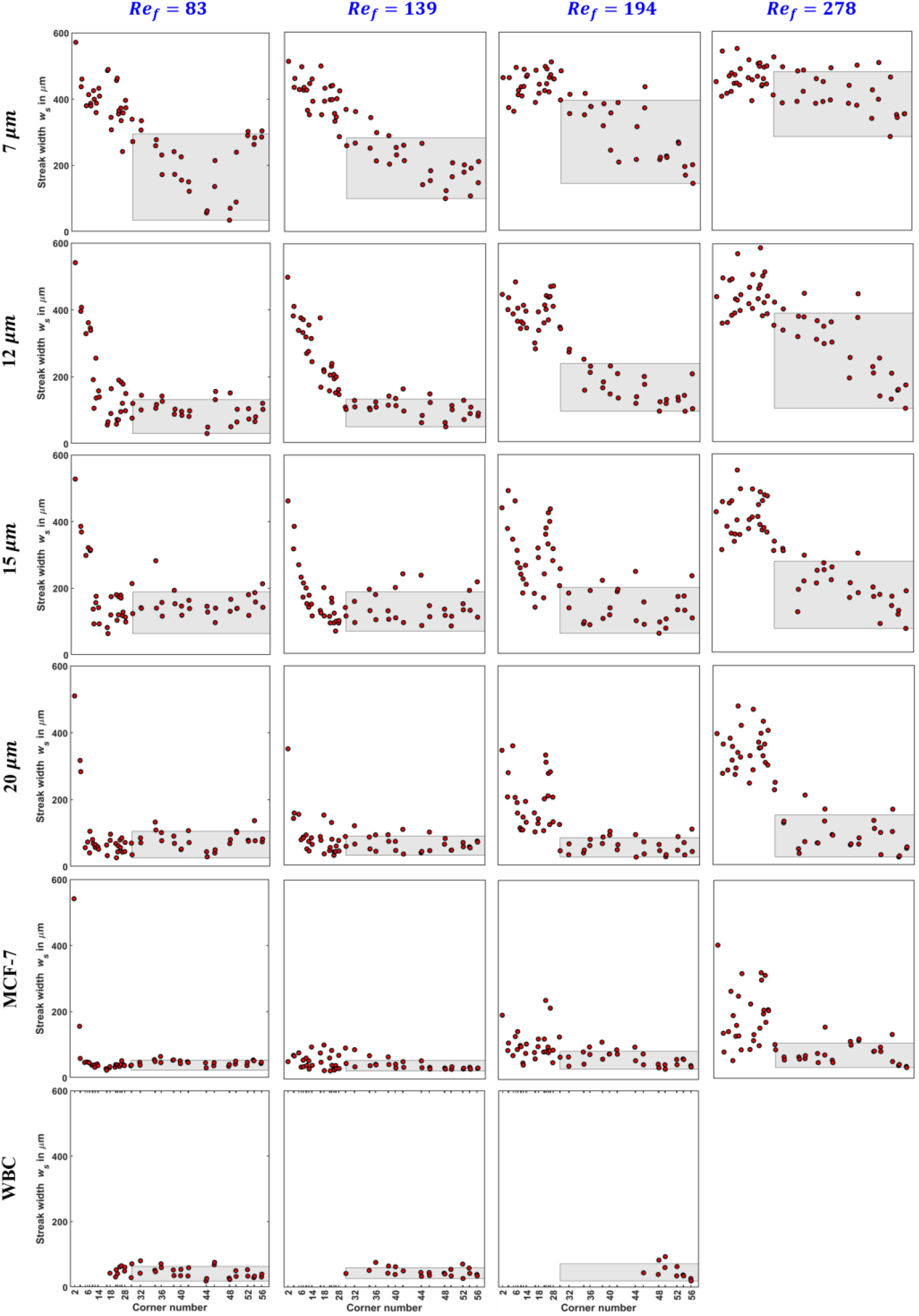
Streak width data for particles and cells at different fluid Reynolds numbers. Streak width *w*_*s*_ is plotted for 7, 12, 15 and 20 *μm* particles as well as MCF-7 breast cancer cells and WBCs at *Re*_*f*_ = 83, 139, 194, 278 respectively. The gray shaded box shows the selected region (ROI) spanning corners 30 to 56 for measuring focusing dispersity at the turns (*FD*_*t*_). The floor is defined by the lowest streak width (*w*_*min*_) among all the labyrinth corners. The roof is defined by *μ* + *σ*, where *μ* and *σ* denote the mean and standard deviation of all the streak widths in the ROI. WBC data at *Re*_*f*_ = 278 is not shown due to weak fluorescence signal resulting from lack of focusing.

To quantify the degree of fluctuations in the streak width data due to turns we develop a measure of focusing dispersity *FD*_*t*_. We chose a region of interest (ROI) between corners 30 and 56 in the labyrinth device, away from the high streak width region. We defined the floor of the shaded ROI as the lowest streak width value *w*_*min*_ among all the corners in the labyrinth. The roof is calculated as: (*μ* + *σ*), where *μ* and *σ* denote the mean and standard deviation of all the streak widths in the ROI. From this, we quantify focusing dispersity as: *FD*_*t*_ = *h*_*s*_/*d*_*p*_, where *h*_*s*_ (= *μ* + *σ* − *w*_*min*_) is the height of the shaded ROI. The ideal case of tight focusing arises when *h*_*s*_ = *d*_*p*_ i.e., *FD*_*t*_ = 1, which would indicate that the fluctuations in *w*_*s*_ are of the same order as particle size *d*_*p*_.

Fig. 3 shows the focusing dispersity for all the conditions investigated. For particles, we observe that for any given *Re*_*f*_, *FD*_*t*_ decreases with increase in particle size. For the 12, 15 and 20 *μm* particles, as well as MCF-7 cells, the focusing dispersity increases at higher *Re*_*f*_ = 278, suggesting greater variability in focusing. Interestingly, WBCs, despite their smaller size (*d*_*p*_ = 10 *μm*), displayed lower dispersity (*FD*_*t*_ = 3 − 6) compared to 15 *μm* particles (*FD*_*t*_ = 8 − 14). Likewise, the MCF-7 cells (*d*_*p*_ = 21 *μm*), have slightly better focusing dispersity (*FD*_*t*_ = 1 − 4) than 20 *μm* rigid particles (*FD*_*t*_ = 2 − 7). Unlike particles, for both cell types, increasing fluid inertia has a less pronounced effect on the measured *FD*_*t*_.

**Figure 3.**
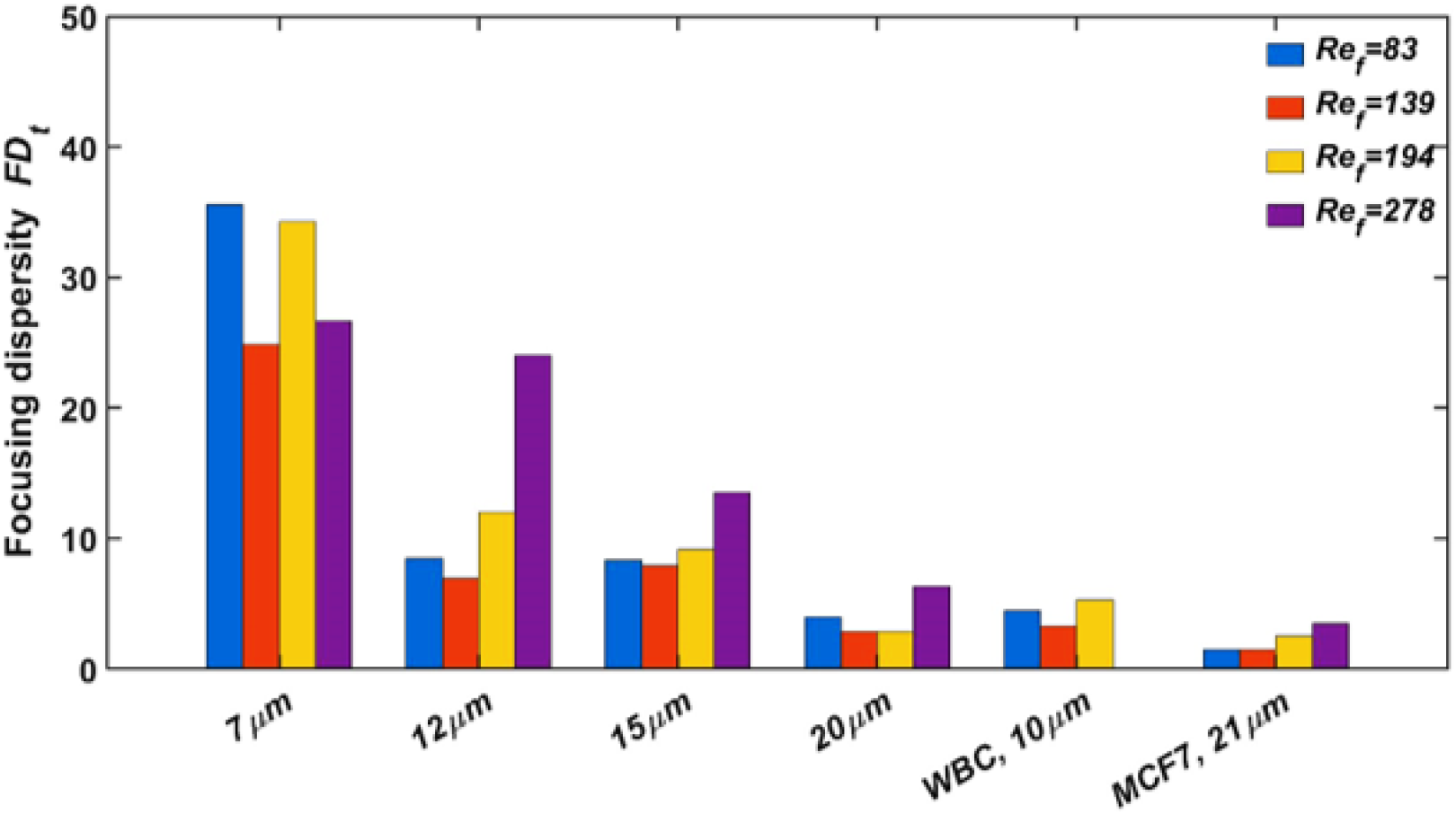
Focusing dispersity of particles and cells in labyrinth device. Focusing dispersity at the turns *FD*_*t*_ is defined as the height of the shaded rectangle in Fig. 3 normalized by the mean size *d*_*p*_. The lower the value of *FD*_*t*_ the less is the variability in the focusing streak width. Results are shown for all *d*_*p*_, *Re*_*f*_ tested.

To explain the effect of particle size, we note that for larger particles, stronger inertial lift forces are able to overcome the cross-sectional mixing forces induced by curvature, resulting in tight streaks without much fluctuation as indicated by the lower values of focusing dispersity ^14,28^. For small particles, the “mixing” forces dominate, causing higher streak dispersion. At high *Re*_*f*_ = 278, the increased strength of curvature forces disturbs the particle streaks even further causing significant variability in focusing. In the case of cells, it seems that tight focusing can be achieved that is less sensitive to *Re*_*f*_, probably due to additional lift forces arising from cell deformability^29^.

### 3.2 Cross-sectional flow visualization in a U-turn using CFD simulations

From the previous section, we observed that in the Labyrinth device as particles are advected downstream, traversing the numerous corners, their streak width measured at the corners fluctuates in a non-monotonic fashion. These fluctuations in streak width could arise due to changes in flow field as the fluid moves from a spiral section into a U-shaped turn and exits into a spiral section. We also evaluated whether the streak width increases or decreases after the entry and exit in the U-turn and did not find any systematic trend suggesting that the turns may not always be beneficial for particle focusing (Fig. S2). In this section, we performed three-dimensional CFD simulations to characterize the flow kinematics at various locations in a U-shaped turn. The observed cross-sectional flow fields at various fluid Reynolds number, help to explain the fluctuations in streak width.

To study the fluid transition behavior in the U-shaped turn, we chose a representative corner pair 55-56, as shown in Fig. 4a. We divided the entire geometry into three regions: (1) entry, containing the spiral section prior to corner 55, (2) straight section between corners 55 and 56 and (3) exit consisting of the spiral segment downstream of corner 56. A total of 8 cross-sectional cuts are taken spanning these regions as shown in Fig. 4a. A qualitative view of how the fluid motion changes in the U-shaped turn is shown in Fig. 4b. The fluid is moving clockwise as it enters the spiral arc and is expected to be pushed towards wall 2. The fluid then changes direction as it traverses through the horizontal section, exits the spiral arc in a counterclockwise direction with the fluid pushed towards wall 1. This change in direction may lead to a cross-sectional flow, the strength of which can depend on the fluids Reynolds number.

**Figure 4.**
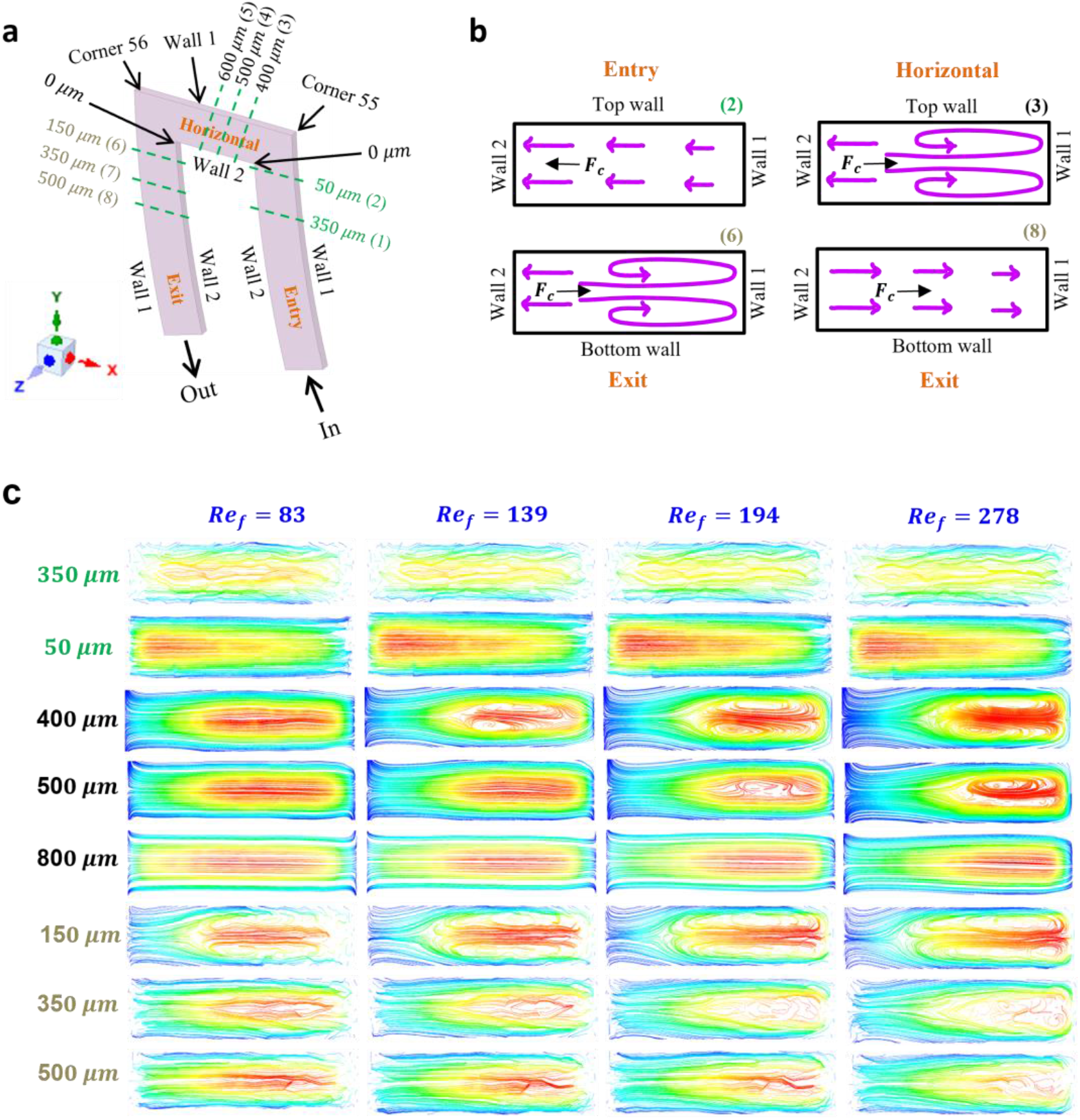
Visualization of cross-sectional flow in the U-turn using CFD simulations. (a) Geometry used for simulations, coordinate axes used are shown on the left. The lateral channel walls have been designated as 1 and 2. This is preferred over the “inner” and “outer” notation as the direction of flow changes from clockwise at the corner entry to counter-clockwise at the horizontal and exit regions. For visualizing the cross-sectional flow, cuts are taken at 8 locations along the entire geometry. Their distances from the respective corner points (0 *μm*) is indicated (not to scale); (b) Schematic showing cross-sectional fluid velocity profiles at 4 selected locations. These are color-coded to match the descriptions in (a). Direction of the centrifugal force *F*_*c*_ is shown by the black arrow; (c) 2D fluid streamlines are displayed at all 8 cross-sectional locations.

Fig. 4c shows the 2D cross-sectional fluid streamlines computed at different downstream distances measured from the corner points, spanning the three regions of interest described above at different *Re*_*f*_. Two features of the flow are evident – a high velocity zone (shown in red) and a vortical flow. We observe that the high velocity zone moves from wall 2 to wall 1 as it enters from the spiral arc into the horizontal section. The vortical flow is most prominent at the higher *Re*_*f*_. We observe the vortices at the entrance of the horizontal section, which appears to vanish mid-way and re-appear at the exit of the spiral arc and disappear downstream.

It is now evident that the presence of sharp turns in the Labyrinth device leads to the generation of Dean vortices. Given that the labyrinth has numerous such turns, it is expected that the local hydrodynamics in the turn plays an important role in dictating the overall focusing performance of the device. Rather than the Dean vortices aiding in particle focusing at the U-turns, we suggest that these vortices are responsible for causing the streak dispersions discussed in the previous section. The ensuing vortex-driven circulatory forces act to disrupt existing equilibrium positions of particles. Larger particles are associated with stronger inertial lift forces which helps them withstand these destabilizing circulatory forces much more than their smaller counterparts 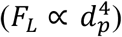. Therefore, their focused streaks are disturbed to a much lower extent.

### 3.3 Effect of U-turn on particle separation

Thus far, we have shown that turns in the Labyrinth device led to streak dispersion at the corners, whose magnitude depends on particle size and fluid inertia. In this section, we evaluate how well separated are the particle focused streams, as it informs on the ability to separate particles of different sizes. To achieve this, we chose the last U-turn (i.e., at corners 55 and 56 where CFD simulations were also performed) in the labyrinth before the downstream expansion section.

We first introduce a measure of effectiveness of particle separation *S* that allows us to assess how well separated are the focused streams of different-sized particles^30^. The higher is the degree of separation, the better is the enrichment of target particles in a mixture. We define the measure of separability 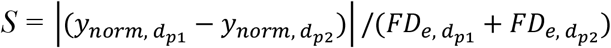, where *y*_*norm*_ is the lateral streak position corresponding to a given particle size, calculated from the peak position of the pixel intensity distribution of the focusing streak, normalized by channel width. As shown in Fig. 5a-i, when two particles are tightly focused and well separated, their separability measure *S* is high. Alternatively, if the peak intensity distributions are overlapping or their focusing tightness is poor, then *S* is low (Fig. 5a-ii,iii).

**Figure 5.**
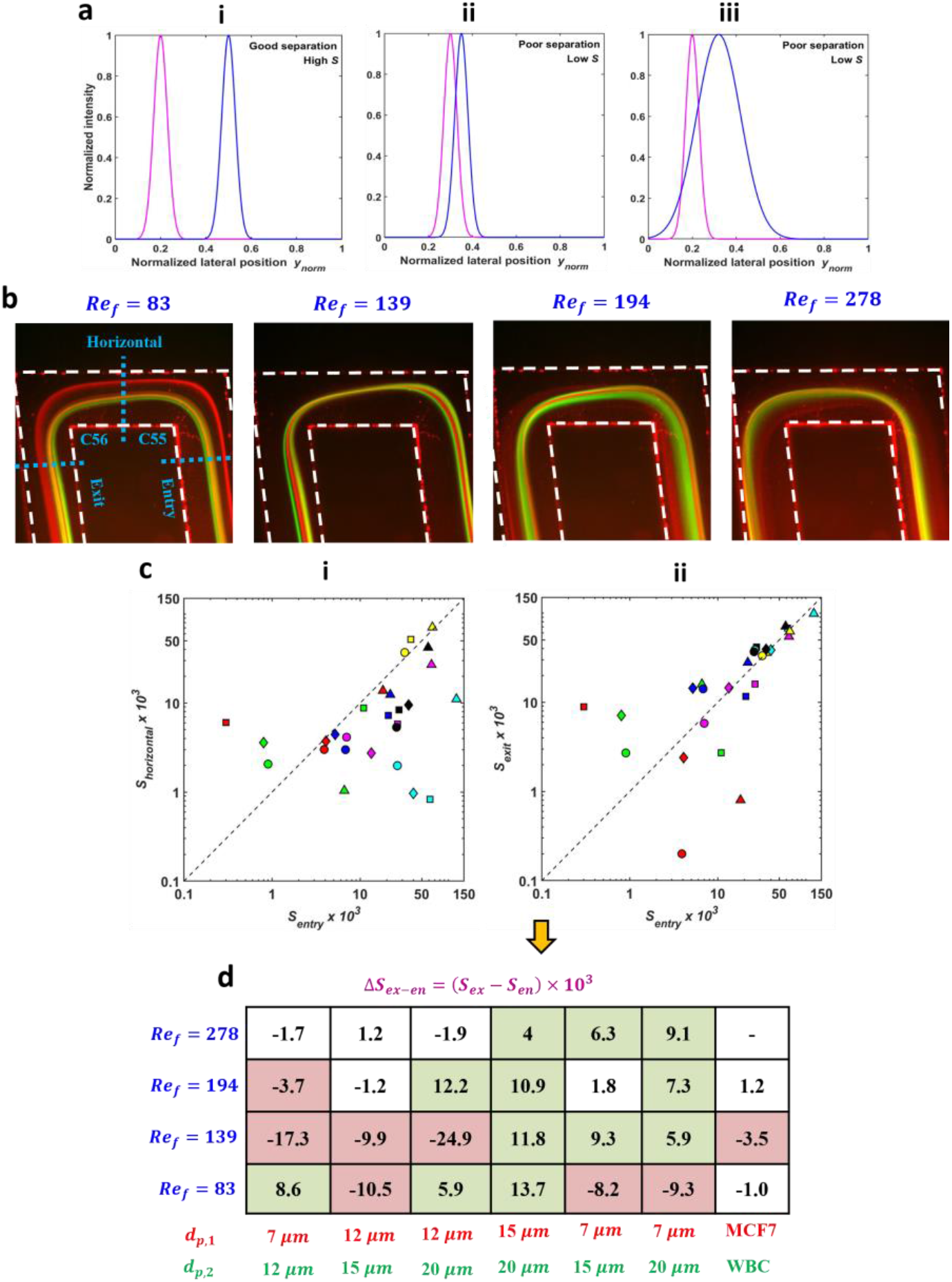
Effect of last U-turn on particle separability. (a) Schematic showing three different cases of separation; (b) Fluorescent streak images of 7 (red), 12 (green), 15 (blue) and 20 (cyan) *μm* particles shown at all *Re*_*f*_ tested. Images are captured near corners 55 and 56 of the Labyrinth device. Channel walls are highlighted by the white dotted line; (c) Separability S is plotted on a log-log scale in the following regions of the Labyrinth device: (i) corner entry and horizontal section and (ii) entry and exit. In each sub-figure, the black dotted line indicates the case where the 2 separability values are identical. Color coding is used to show the different size pairings: 7,12 *μm* (red), 12, 15 *μm* (green), 12, 20 *μm* (blue), 15, 20 *μm* (magenta), 7, 15 *μm* (cyan), 7, 20 *μm* (black) and MCF7, WBC (yellow) respectively. Marker shape is used to illustrate *Re*_*f*_: 83 (square), 139 (triangle), 194 (round) and 278 (diamond); (d) Table shows the difference in the pairwise separability values Δ*S*_*ex*−*en*_ between the corner entry and exit regions for all particle and cell sizes (*d*_*p*_) and *Ref* tested. Cases corresponding to positive and negative changes are shaded in green and red respectively. Cases where Δ*Sex*−*en* is lower than the measurement limit are not colored. Cell data is not shown at *Re*_*f*_ = 278 as WBCs did not focus and therefore separate at this condition.

Using the separability measure, we seek to evaluate the effect of the sharp turn on the separation between size-varying particles. Two important questions worth pursuing are: (1) how does the separability change as particles moves into the horizontal section of the U-turn and (2) after exiting the turn, does the separability improve or deteriorate.

Fluorescence streak images of 7, 12, 15 and 20 *μm* particles at various flow conditions are first captured in the turn region. Fig. 5b shows composite RGB images of the particle streaks for all four flow conditions tested. To effectively capture the separation in the turn region, we divide the geometry into three segments: (1) entry – spiral segment preceding corner 55, (2) transition – straight section between the two successive corners and (3) exit – spiral segment succeeding corner 56 (Fig. 5b). This division is helpful because it allows us to contrast particle separability before and after the turn, enabling us to gauge its impact on separation.

In these three regions, we compute pairwise separability as before. The 2D scatter plot in Fig. 5c shows the separability S computed in two of the above-mentioned regions for all particle sizes and *Re*_*f*_ tested. A comparison of S at corner entry and transition shows that in general, S decreases adversely in the latter region (Fig. 5c-i). This behavior may be attributed to the tight curvature of the turn. Particles traversing the radially lenient spiral curve are suddenly subjected to very strong turn curvature. This results in a notable increase in the inertia-driven centrifugal forces which leads to lateral crisscrossing of particle streaks. As an immediate consequence, the well separated streaks in the preceding spiral section now become almost indistinguishable. The only major exception here occurs in the case of 7 and 12 *μm* particles at *Re*_*f*_ = 83. Here, due to insufficient inertia, we do not observe lateral streak movement and their motion follows the geometry of the turn, preserving separation.

Next, we compare separability in the two spiral sections before (entry) and after (exit) the turn to understand its overall effect on particle separation (Fig. 5c-ii). Interestingly, it is found that turns may improve or deteriorate particle separation depending on their sizes and strength of fluid inertia. The table in Fig. 5d reports the difference in separability between the entry and exit regions measured as Δ*S*_*ex*−*en*_ = (*S*_*ex*_ − *S*_*en*_) × 10^3^. To quantify the level of measurement noise, we obtained Δ*S*_*ex*−*en*_ over three independent trials and found the standard deviation to be about 2 consistently over all *Re*_*f*_ tested. Positive values (in green) indicate improved separation due to the U-turn and negative values (in red) indicate poor separation. Based on the data, it is clear that U-turns can be beneficial or detrimental depending on the particle size in the mixture, and the fluid Reynolds number, although it appears that for the 15 and 20 *μm* particle combination, U-turn might be beneficial. Compared to similar-sized particles, the separation between MCF-7 cells (*d*_*p*_ = 21 *μm*) and WBCs (*d*_*p*_ = 10 *μm*) is almost unaffected by the sharp turn.

### 3.4 Effectiveness of separation at the expansion section in the labyrinth

Finally, we apply our developed measure of separability in the downstream expansion region (Fig. 7a) to quantitatively evaluate how well labyrinth is able to separate particles and cells of different sizes. Even though the labyrinth has multiple U-turns, ultimately, the enriched cell/particle populations have to traverse the expansion section where the separability could be altered. Since these measurements are carried out in the expansion near the device exit, where the purified populations may be collected from independent outlets, the separability measure here serves as an overall performance indicator of size-based enrichment.

Fig. 7b shows representative overlay images for two different target sizes and their separability scores. We find that the fluorescent particle streaks for the 15 and 20 *μm* at *Re*_*f*_ = 194, nearly overlay, giving a low value of S = 0.0013 (Fig. 7b-i). For the 7 and 12 *μm* at *Re*_*f*_ = 83, the fluorescent steaks are a bit more separated, giving S = 0.0064 (Fig. 7b-ii). For the MCF-7 and WBC at *Re*_*f*_ = 139, the separability S = 0.0450 is high due to the widely separated fluorescent streaks (Fig. 7b-iii).

Next, we show pair-wise comparisons for all conditions investigated in Fig. 7c. Among particles, the best separation is obtained between 12 and 20 *μm* at *Re*_*f*_ = 83 (S = 0.0137). At the same flow condition, 12 and 15 *μm* particles can also be adequately separated (S = 0.0078). In both cases, increasing the flow rate results in a non-monotonic decrease in separability. Additionally, 7 *μm* particles may be sufficiently separated from 12 *μm* at *Re*_*f*_ = 194 (S = 0.0066). However, they cannot be separated to a similar degree from 15 and 20 *μm* particles with S = 0.0035 and 0.0046 respectively. We also find that 15 and 20 *μm* particles are not able to be adequately separated at any *Re*_*f*_ in spite of their size differences. Finally, at the highest flow rate tested (*Re*_*f*_ = 278), none of the particles could be separated based on size (S < 0.003). From these results, we infer that there is no unique *Re*_*f*_ that yields effective separation of all particle sizes tested. Consequently, at a single flow condition, attempts to separate more than two particle sizes would result in a significant loss of separation purity.

Even though the separation performance of labyrinth on rigid particles is poor, it is effective at separating cancer and white blood cells. At the lower flow rates (*Re*_*f*_ = 83, 139), we find that labyrinth was able to separate MCF-7 cells from WBCs extremely well with S = 0.0256 and 0.0450 respectively. This is significantly higher than what was observed with particles, suggesting that size alone may not be able to account for the enhanced separation in case of cells. The promising separation achieved did not however, hold at higher *Re*_*f*_, wherein S < 0.001 and these cells could not be separated.

Next, we assess whether the separability at the exit of the last turn in labyrinth (Fig. 5c-ii) is conserved at the downstream expansion section. The linearly tapered expansion section is usually considered to magnify the spacing between particle focused streams, however, it also reduces inertial force on the particles potentially affecting the separability. Fig. 6d shows the separability S measured in these two regions for all sizes and flow conditions tested. A striking observation here is that the separability significantly reduces in the expansion as compared to the turn exit. Although an incremental increase in the channel width in the taper helps to improve the peak-to-peak spacing of particle streams, it is found that the focused streaks are relatively diffuse in this region which lowers separability. We suggest that the divergent nature of the expansion reduces the velocity of the advecting fluid (*U*_*f*_). Consequently, the inertial lift force responsible for focusing particles is weakened 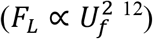 which results in streak widening and therefore lowers the separability compared to the turn exit.

**Figure 6.**
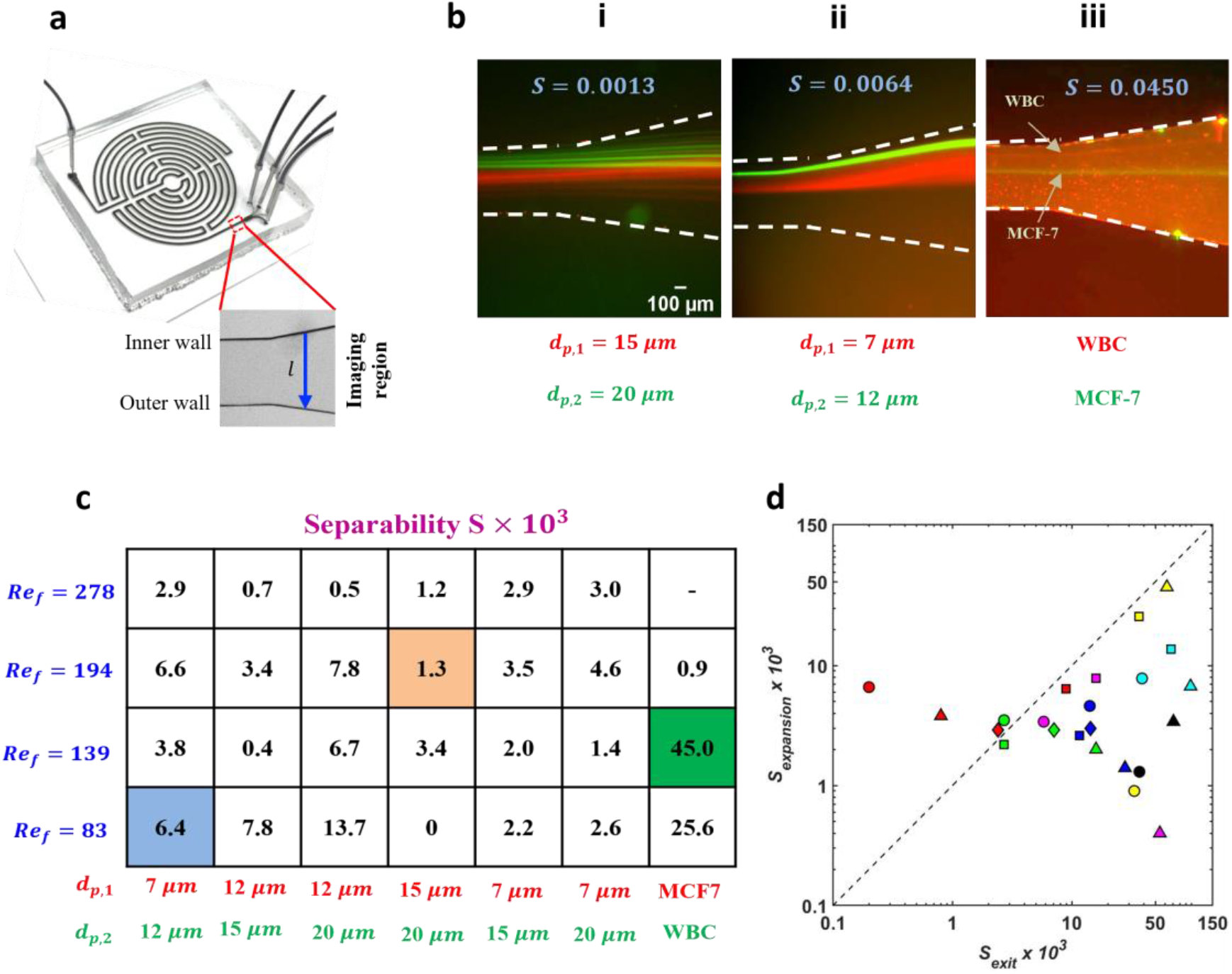
Separation performance of labyrinth in the expansion section. (a) Measurements are performed in the expansion region (inset) of the labyrinth device in the direction of the blue arrow; (b) Three representative cases of low, medium and high S is shown by creating a composite image between the two particle sizes or cell types tested; (c) Separability S between a pair of particle sizes in the downstream expansion region. To obtain S, each value in the table is to be multiplied by 10^−3^. The colored squares indicate S values corresponding to the images in (b); (d) 2D scatter plot showing separability S at the exit of the last turn (*S*_*exit*_) and downstream expansion region (*S*_*expansion*_). The dotted line represents the situation when *S*_*expansion*_ = *S*_*exit*_. Marker shape and color descriptions are the same as Fig. 5c.

## 4. Conclusion

In this work, inertial, size-dependent focusing was investigated in the microfluidic labyrinth device, whose geometry consists of a multitude of sharp turns along with circular loops. Particularly, we investigated whether the addition of sharp curvatures helped to increase separation by virtue of inducing strong circulatory flows in the cross-section. First, we characterized the focusing dynamics inside the device by testing particles and cells of various sizes (*d*_*p*_) at different fluid Reynolds numbers (*Re*_*f*_) and measured the width of the fluorescent streaks at all 56 corners. At all the conditions, we observed intermittent fluctuations, wherein particles that initially focused at a corner defocused at other downstream corners. To quantify these streak dispersions at the turns, we devised a measure, focusing dispersity *FD*_*t*_. In general, it was seen that *FD*_*t*_ was lower for larger particle sizes and at lower *Re*_*f*_ indicating better focusing. Moreover, the focused streaks of cells were found to be much more immune to changes in *Re*_*f*_.

To test our hypothesis that the streak fluctuations stem from variations in the cross-sectional flow-field induced by the complex corner geometry, we conducted 3D CFD fluid simulations. Our results confirmed the presence of prominent Dean vortices located at specific positions downstream of the corner. Induced by the sharp curvature of the turn, we suggest that this vortical flow acts to disturb the equilibrium positions of particles and is responsible for the for the observed streak fluctuations at the corners.

Next, we developed a measure separability to determine how the separation varies as particles traverse a sharp turn. We found that the turn does not always improve particle separability and positive and negative outcomes depend on *d*_*p*_ and *Re*_*f*_ tested. Furthermore, we observed that S substantially varies along the turn which can be attributed to the presence of the afore-mentioned corner-induced Dean vortices.

Finally, we measured separability at the device exit to evaluate how well labyrinth is able to enrich particles or cells of a given size, impacting purity. We found that there was no unique flow condition where all four tested sizes could be efficiently separated and a careful choice of the optimal *Re*_*f*_ would have to be made depending on the sizes present in the heterogeneous mixture. Additionally, we found that cells, unlike particles of similar sizes, could be separated to a much higher degree as evidenced by the order of magnitude increase in their separability values.

In summary, our work is one of the first efforts to study inertial focusing dynamics of particles and cells in curved geometries involving sharp turns. Our results in the labyrinth device reveal that turns can either improve or reduce particle separation depending on their sizes and flow conditions tested. Cells on the other hand, were found to be relatively much more resistant to the sharp turns. Further targeted studies are needed in future for a more mechanistic understanding of these findings.

## 5. Author contributions

AG designed and ran the experiments, performed the analysis and generated all the data. AG and SAV wrote the paper, SAV provided feedback on the manuscript and was also the supervisor of the project.

## 6. Acknowledgements

The authors would like to thank members of the Nagrath laboratory for providing the Labyrinth devices to us and are grateful to them for useful discussions. We would like to acknowledge Guanqiao Feng, PhD for helping us develop the separability measure used in this study. This work was funded by the Cancer Prevention Research Institute of Texas (CPRIT, Grant no. RP190658).

## 9. Supplementary Information

**Figure S1.**
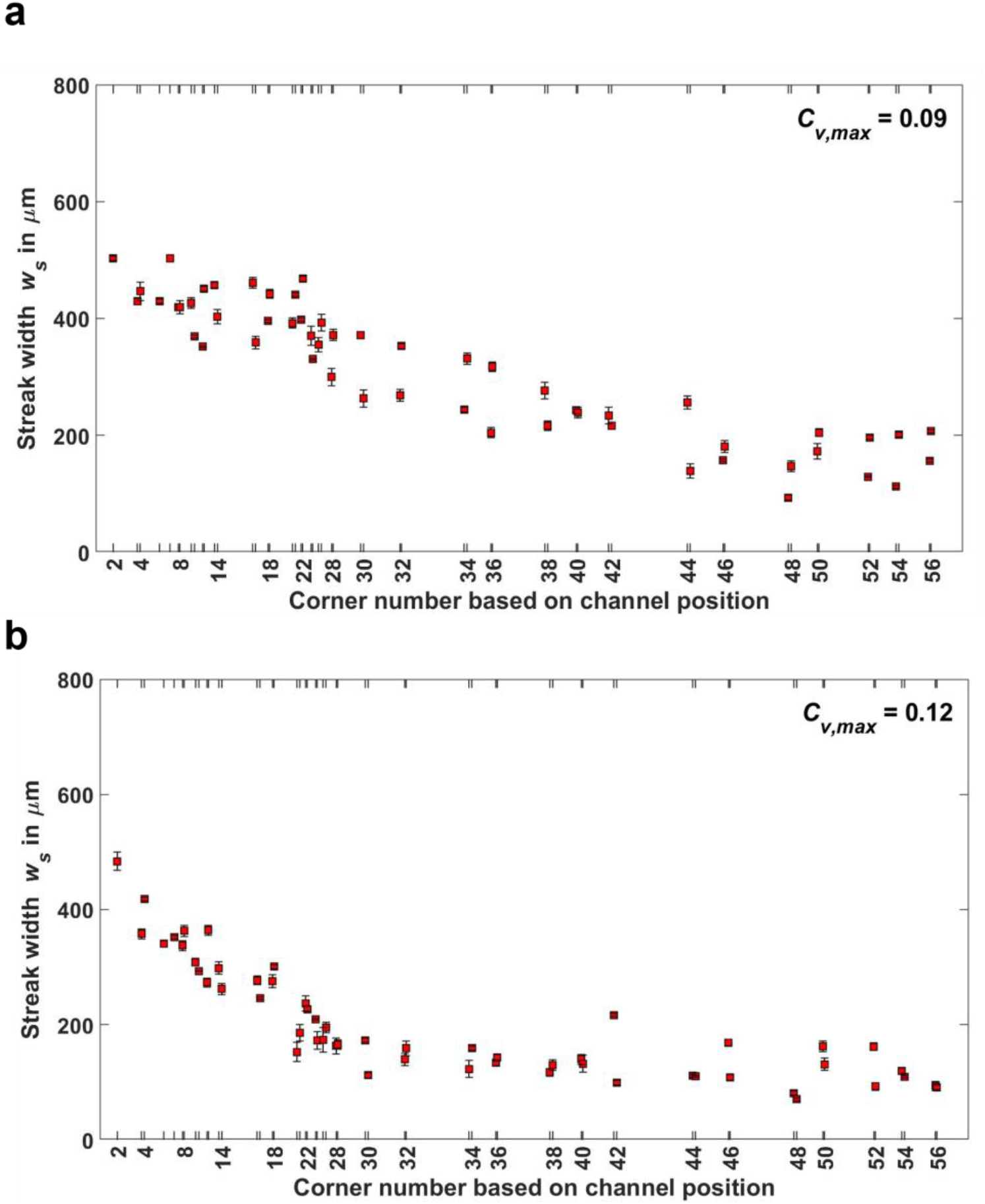
Repeatability of *w*_*s*_ measurements. Streak width measurements *w*_*s*_ are shown at each of the 56 corners for 7 (a), 12 (b) *μm* particles at *Re*_*f*_ = 139. Each data point indicates the average of three replicate measurements, error bars show the standard deviation. Among all corners, maximum coefficient of variation *C*_*v, max*_ is calculated as 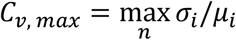, where σ_*i*_, *μ*_*i*_ denotes mean and standard deviation for each corner point *i, n* is the total number of corner points (*n* = 55).

**Figure S2.**
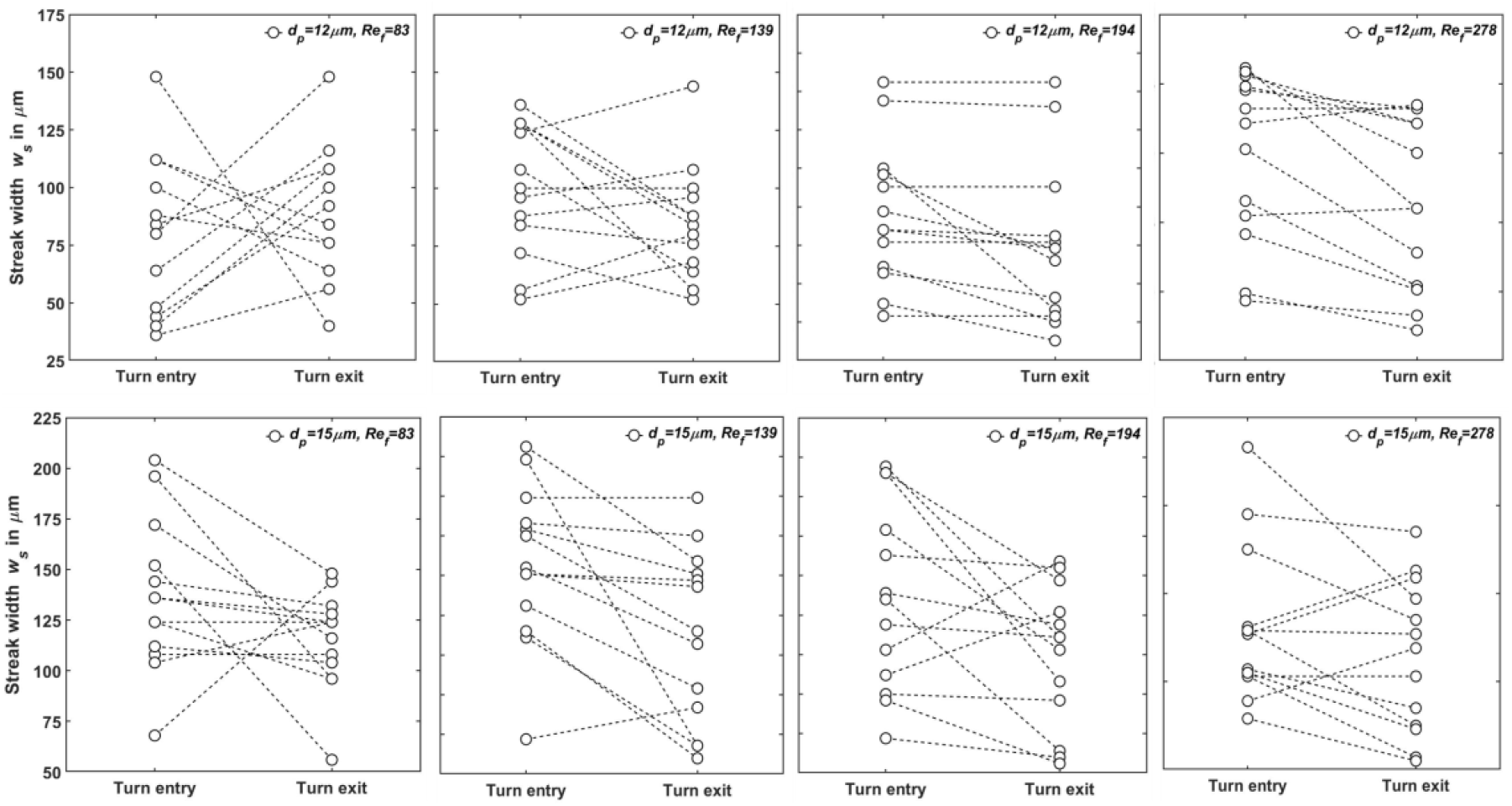
Streak width variation from corner entry to exit. Streak widths measured at the turn entry and exit regions are shown for 12 and 15 *μm* particles at all *Re*_*f*_ tested. The dotted line bounded by the two circles at the ends represents a single turn, data is shown for 13 such turns. To capture the effect of the turn on focused streaks, only turns after corner 30 are considered, this ROI is consistent with that reported in Sec. 3.1.

**Figure S3.**
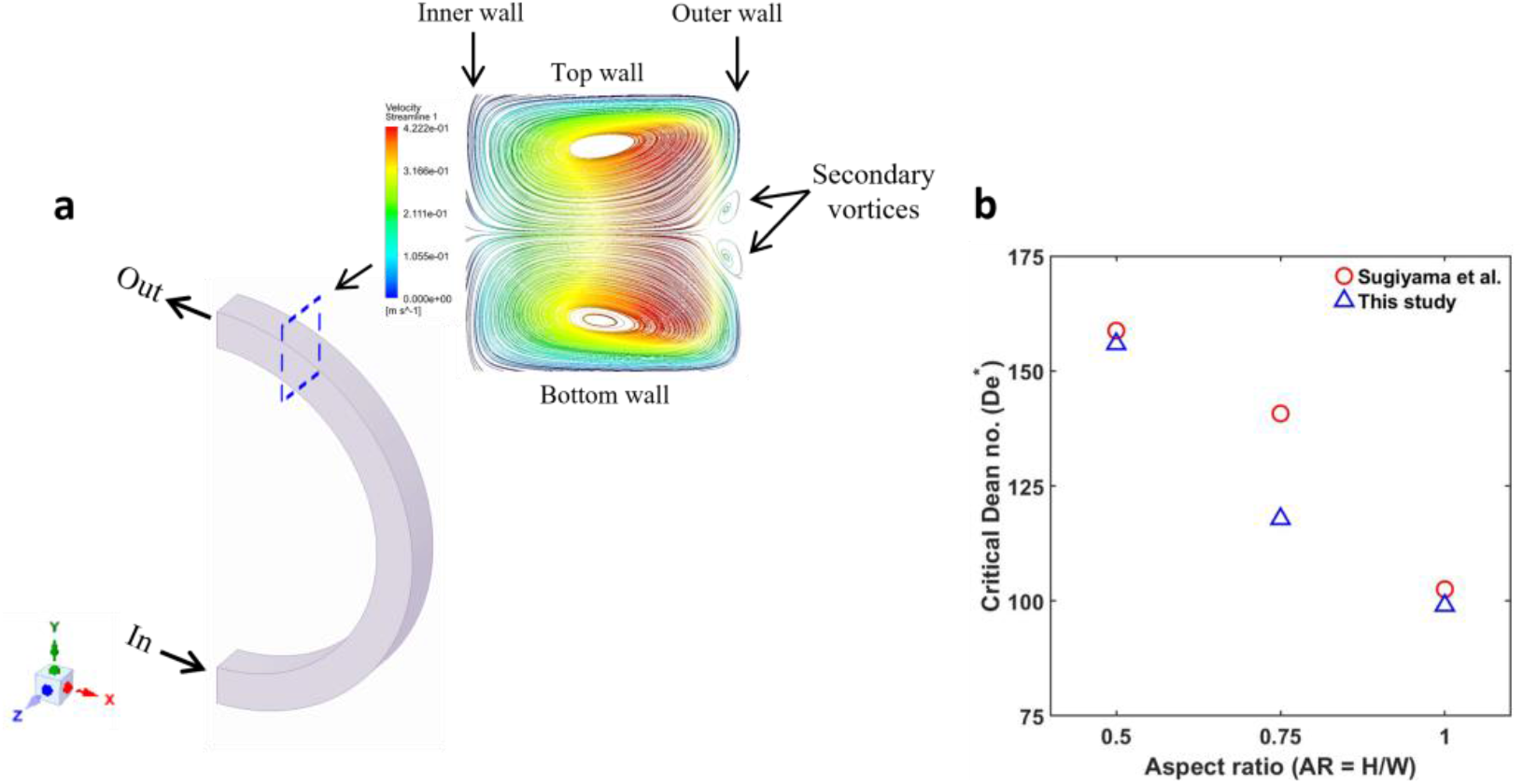
Benchmarking CFD simulations. (a) Curvilinear geometry used for simulations. Inset shows a snapshot of the cross-sectional flow streamlines at the point of transition for aspect ratio of 1. Two additional smaller fluid vortices appear near the outer wall; (b) The corresponding critical Dean number *De*^*^ is reported for three different cases of channel aspect ratios tested. Results are benchmarked against the experimental data reported by Sugiyama *et al*.

We benchmarked our CFD simulations, by selecting a curvilinear geometry where the curvature ratio *δ* varies from 0.067 to 0.1 and the aspect ratio (*AR* = *H*/*W*) increases from 0.5 to 1. We selected the study by Sugiyama *et al*. ^31^, who reported the influence of channel aspect ratio on the onset of secondary Dean vortices characterized by the critical Dean number *De*^*^. This was defined as the Dean number (*De*) at which the two primary cross-sectional vortices split into four (Fig. S3 a). They performed experiments and showed that *De*^*^ decreases with increase in the channel aspect ratio (Fig. S3 b) but the effect of curvature ratio is relatively less pronounced. To obtain this data from our simulations, we iteratively increased *De* by virtue of the imposed flow rate and tracked the onset of transition. Generally, we obtain good agreement and expect our benchmarking to be applicable to the labyrinth geometry, which has an aspect ratio of 0.2.

